# FIRST RECORD OF *MELANAPHIS SORGHI* (THEOBALD, 1904) (HEMIPTERA APHIDIDAE) IN ITALY AND SPAIN

**DOI:** 10.1101/2023.05.02.539111

**Authors:** Alice Casiraghi, Nicola Addelfio, Nicola M. G. Ardenghi, Nicolás Pérez Hidalgo

**Affiliations:** Instituto Valenciano de Investigaciones Agrarias. (IVIA). Unidad de Entomología, Centro de Protección Vegetal y Biotecnología. Ctra. Moncada-Náquera Km. 4,5. E-46113 Moncada, Valencia, Spain; Ministero della Cultura, Gallerie degli Uffizi, Piazzale degli Uffizi 6, 50122 Firenze (FI); Botanic Garden, University Museum System, University of Pavia, via S. Epifanio 14, I-27100, Pavia, Italy; Departamento de Artrópodos, Museo de Ciencias Naturales de Barcelona, 08003, Barcelona, Spain

**Keywords:** *Sorghum* aphid, alien species, pest, Italy, Iberian Peninsula

## Abstract

The sorghum aphid *Melanaphis sorghi* (Theobald) is recorded for the first time in mainland Italy (Florence, Tuscany region, Italy) and Spain (Vinalesa, Valencia Region, Spain) on *Sorghum halepense* (L.) Pers. Data on its biology, worldwide distribution and notes on its pest behaviour are given. *Melanaphis sorghi* had previously been recorded from Greece (in 2008), Cyprus and Israel. The records of this species in Iran and Turkey need confirmation.

## INTRODUCTION

The genus *Melanaphis* van der Goot, 1917 contains around 25 species (Favret, 2022; Blackman & Eastop, 2022) of palearctic distribution, associated with Poaceae and only a few shows alternating host behaviour between Pyroideae (their primary host) and Poaceae (the secondary host) (Holman, 2009; Blackman & Eastop, 2022). Most of the species of the genus have their origin in East Asia and are associated with *Miscanthus* or *Arundinaria* and bamboos. Four species, *Melanaphis donacis* (Passerini, 1862), *Melanaphis elizabethae* (Ossiannilsson, 1967), *Melanaphis luzullella* (Hille Ris Lambers, 1947) and *Melanaphis pyraria* (Passerini, 1862), are present on European territories (Holman, 2009). Two others, originated in Southeast Asia, *Melanaphis bambusae* (Fullaway, 1910) and *Melanaphis sorghi* (Theobald, 1904), have been introduced in Europe (Mediterranean basin and south) and other parts of the world (Ortego et al., 2004; Holman, 2009; Margaritopoulos et al., 2013; Nibouche et al., 2021; Balbi et al., 2022). However, the genera needs a deep revision combining biological aspects with morphological and molecular data (Nibouche et al., 2021; Blackman & Eastop, 2022). In Italy and Spain three of them (*M. pyraria, M. donacis, M. bambusae*) are well known and are widely distributed on both peninsulas (Pérez Hidalgo & Mier Durante, 2005; Barbagallo et al., 2011).

Citizen science has proven to be a very powerful tool for monitoring and contributing to new data on biodiversity from every corner of the world. Social networks such as Facebook (in its groups dedicated to gardening or nature lovers), iNaturalist or Biodiversidad Virtual (mainly in Spain) have allowed the identification of new records of invasive species (Chamberlain, 2018; Jaskuła et al., 2022).

iNaturalist database, since its creation in 2008, has accumulated more than 125 million observations (photos and audio recordings) and each day receives an average contribution of 60,000 new items. It represents an important source of information on records, ecology, species distribution, especially regarding invasive species or those that are vulnerable and difficult to find (Hiller & Haelewaters, 2019; Serniak et al., 2022). An observation accompanied by a photograph of an aphid colony taken on November 4^th^, 2022, in Florence and uploaded to the iNaturalist.org platform (https://www.inaturalist.org/observations/141154007) by the second of the authors, has made it possible the identification of an invasive Aphididae.

## MATERIALS AND METHODS

Material studied form Italy. A sample (reference 4417) collected on November 8^th^, 2022, on *Sorghum halepense* (L.) Pers. along the Terzolle and Mugnone streams in Florence (coordinates taken from Google.maps: 43.790094, 11.232194) that was composed by several apterous and winged viviparous females (November 13^th^) and were probably attended by workers of *Tapinoma nigerrimum* group.

Material studied from Spain. A sample (reference 4474) with many apterous viviparous females recorded on *S. halepense* on January 15^th^, 2023, in Vinalesa (Valencia) (coordinates taken from Google.maps: 39.542472, -0.373399).

Apterous viviparous females from Italy and Spain were prepared and measured (25 from the Italian sample and 19 from the Spanish sample) to confirm the identification of the aphid species following the ID keys of Blackman & Eastop (2022). The samples in ethanol and all slides with the measured specimens are deposited in the aphidological collection of the “Instituto Valenciano de Investigaciones Agrarias” (Valencia, Spain). The host plant was identified by Nicola M. G. Ardenghi.

## RESULTS AND DISCUSSION

The morphological and molecular studies carried out by Nibouche et al. (2021) allow a morphological separation of *M. sorghi* and *M. sacchari* based on the ratio between the length tibia of the hind legs and the processus terminalis of the last antennal segment: 1.8-3.0 for *M. sorghi* and 1.5-1.9 for *M. sacchari*.

The identification of the samples by the keys of Blackman & Eastop (2022) allowed us, without doubts, confirm the presence of *M. sorghi* for the first time in mainland Italy and Iberian Peninsula (Spain). The Italian apterous viviparous females (n= 25) present a ratio of 1.82 to 2.44, with mean 2.063 and the Spanish (n= 19) ones 2.17 to 2.66, mean 2.35.

Apterae of *M. sorghi* are white or yellow (Fig. 1A, 1B, 1C) and sometimes pinkish with siphunculi black. Some larger individuals have a variably developed of black dorsal abdominal patch (see also influentialpoints.com/Gallery). Alatae (Fig. 1D) are similar to the apterae but with more secondary rhinaria on antennal segments (III with 4-13, IV with 0-1 and V with 0). Nibouche et al. (2021) gives very good descriptions and illustrations of all the forms and characteristics that allow its separation from *Melanaphis sacchari*.

**Figure 1.**
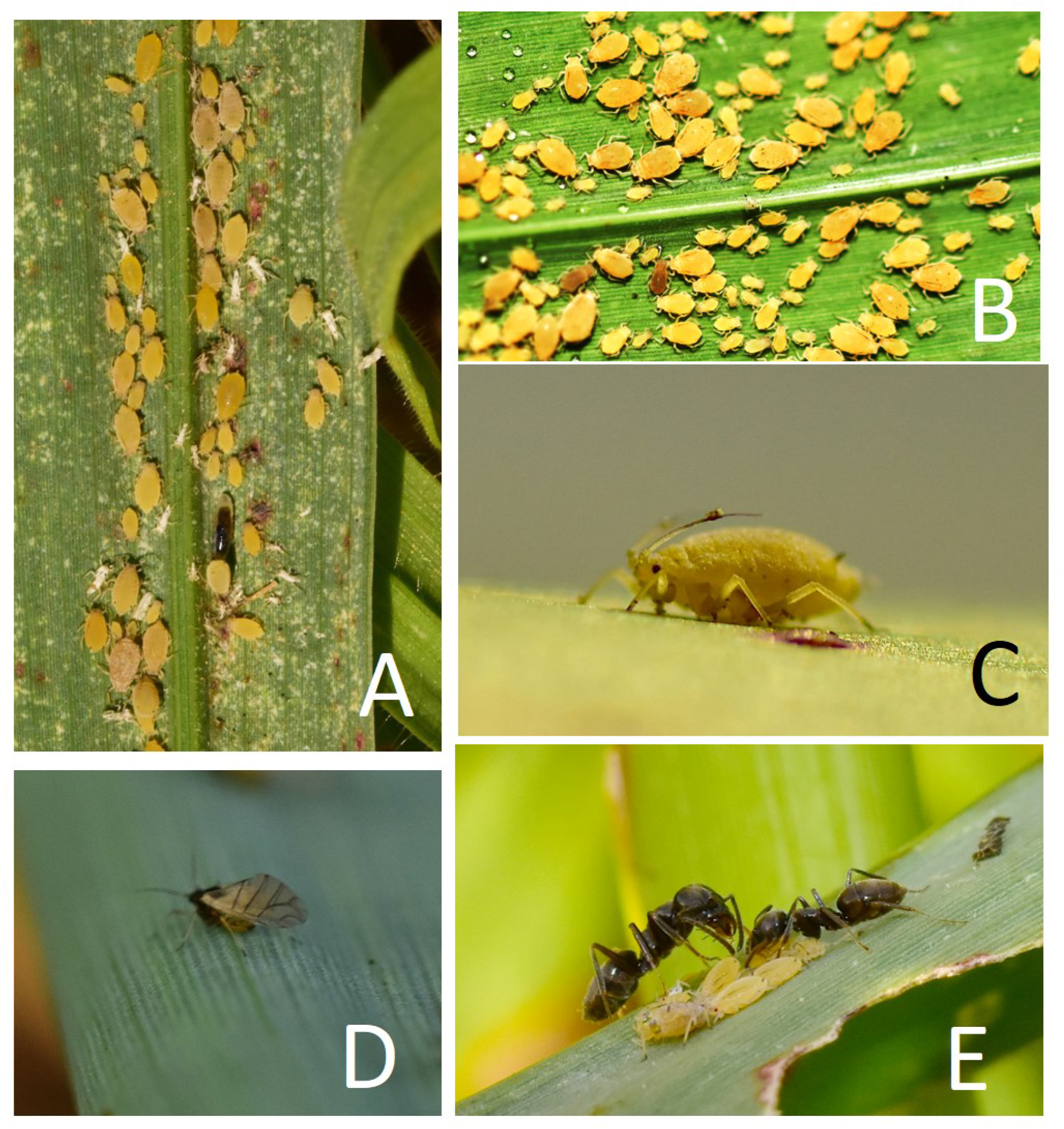
Different colonies of *Melanaphis sorghi* on *Sorghum halepense* in Florence (Italy) (A, B), apterous viviparous female (C), winged viviparous female (D) and two workers of *Tapinoma nigerrimum* group tending several nymphs (E).

*Melanaphis sorghi* lives on *Sorghum* taxa [*S. bicolor* (L.) Moench subsp. *bicolor, S. bicolor* subsp. *arundinaceum* (Desv.) de Wet & J.R. Harlan ex Davidse, *S. halepense, S. ×almum* Parodi], *Eleusine* [*E. coracana* (L.) Asch. & Graebn. s.l., *E. indica* (L.) Gaeretn.] and there are also records on *Saccharum officinarum* L., *Cenchrus americanus* (L.) Morrone subsp. *americanus* and *Zea mays* L. subsp. *mays* (Nibouche et al., 2021; Blackman & Eastop, 2022).

The majority of populations of *M. sorghi* appear to be anholocyclic, but in Japan it may behave holocyclically (Setokuchi, 1975) and oviparous have been collected in north-western India in February and March (David, 1977). In addition, in Mexico sexual morphs have been recorded in introduced populations of the *M. sacchari*/*sorghi* group but their belonging to *M. sacchari* or *M. sorghi* has not been confirmed (Peña Martinez et al., 2016).

For a long time, the *Melanaphis sacchari*/*sorghi* complex has been recorded from many parts of the world and on many grass species. For this reason, Blackman & Eastop (2022) indicate that many of the records of this complex on *Sorghum* cultivated in Africa and Asia probably are *M. sorghi*. Until the molecular and morphological studies carry out by Nibouche et al. (2021), it has not been possible to separate the two species. Currently we can ensure that *M. sorghi* is present in east Asia (India, Pakistan, China, Thailand, Japan, Philippines) probably its original range, and has been introduced in Africa (in the south and the west, and in the east only in Kenya and Uganda), in North America (in 17 States of USA and Mexico), South America (Brazil and Argentina) and in different Caribbean countries (Haiti and Puerto Rico) (Nibouche et al., 2014; Nibouche et al., 2021; Harris-Shultz et al., 2022; Balbi et al., 2022).

In Europe, it has been cited for the first time in Greece (in Messolonghi and Thessaloniki; Margaritopoulos et al., 2013) and Cyprus (unknown locality; Blackman & Estop, 2022), and there are also records from Israel (unknown locality; Blackman & Eastop, 2022). The records from Iran (unknown locality; Eastop & Hodjat, 1993, as *M. sacchari*) and Turkey (Adana and Antalya; Köl & Özdemir, 2021, as *M. sacchari*) need to be confirmed because Blackman & Eastop (2022) considered them as *M. sorghi*.

In any case, *Melanaphis sorghi* and *M. sacchari* can live on the same host plants and currently coexist in many territories, therefore many of the records attributed to *Melanaphis sacchari* should be confirmed in light of new taxonomic and molecular knowledge.

This species has become the number one pest of cultivated sorghum in southern USA and Brazil. It produces damage similar to that of another well-known pest in sorghum, *Schizaphis graminum* Rondani, 1852: foliar chlorosis, reduced grain yield, and even the death of the plant. In addition, unlike *S. graminum, M. sorghi* is capable of feeding on the plant during the milky stage of the grain, reducing its quality and allowing the development of bold in the area (Michaud, 2017).

In Italy, sorghum has become an important crop in recent years, reaching 35,375 hectares in 2022, with a production of 1,837,956 quintals in the last year (ISTAT, 2022). It is one of the four largest producers in Europe (Sorghum-ID, 2022), which Emilia-Romagna (19,383 ha), Veneto (4,479 ha) and Tuscany (2,040 ha) as the leading regions. (ISTAT, 2022).

In Spain, sorghum is mainly cultivated in Western Andalusia and Southern Pyrenees. More than 25.000 t were produced in 2020 in the country (Sorghum-ID, 2022). The presence of this invasive aphid could slow down or stop the production of this new cereal in the Mediterranean basin.

The discovery of this new pest on *Sorghum halepense* is further worrisome: in Europe *S. halepense* is an invasive archeophyte (originated from Western Asia) and a widespread noxious weed (especially of maize and soybean), already known as a reservoir for crop pathogens. Its spread is enhanced by a combination of seed and vegetative (through rhizomes) reproduction and its persistence is guaranteed by an increasing resistance to herbicides (Paterson et al., 2020; Follak & Essl, 2012). With these features, *S. halepense* may rapidly become an ally of *Melanaphis* in its expansion across the European continent.

## CONCLUSIONS

It is possible that the sorghum aphid is more widely distributed than what is reflected in this article, both in Italy and Spain and in the rest of Europe, given that sorghum cultivation is increasing in European and North African territories and its favourite host plant, *S. halepense*, is widely distributed.

*Melanaphis sorghi* is one of the species that is currently expanding throughout the world, clearly associated with the movement and cultivation of its main host plant, sorghum. That is why it is not surprising records in the countries of the Mediterranean basin and that citizen science, together with pest control services, help to detect its presence in the near future.

## ACKNOWLEDGEMENTS

The authors to thank Doc. Francesca Maria Gatti (Université de Strasbourg, Master in Translation, specialized in English and Russian) for reviewing the article in English and Prof. Xavier Espadaler (Universidad Autónoma de Barcelona, Spain) for confirming the identification of the ant species group.

